# AMPK signaling linked to the schizophrenia-associated 1q21.1 deletion is required for neuronal and sleep maintenance

**DOI:** 10.1101/391722

**Authors:** Stanislav Nagy, Gianna W. Maurer, Julie L. Hentze, Shu Kondo, Morten Rose, Thomas M. Werge, Kim Rewitz

## Abstract

The human 1q21.1 deletion of ten genes is associated with increased risk schizophrenia. The deletion involves the ß-subunit of the AMP-activated protein kinase (AMPK) complex, a key energy sensor in the cell. Although neurons have a high demand for energy and low capacity to store nutrients, the role of AMPK in neuronal physiology is poorly defined. Here we show that AMPK is important in the nervous system for maintaining neuronal integrity and for stress survival and longevity in *Drosophila.* To understand its impact on behavior and potential contribution to the 1q21.1 deletion syndrome, we focused on sleep, as its main role is proposed to be to reestablish neuronal energy levels that are diminished during energy-demanding wakefulness. Sleep disturbances are one of the most common problems affecting individuals with psychiatric disorders. We show that AMPK is required to maintain proper sleep architecture and for sleep recovery following sleep deprivation. Neuronal AMPKß loss specifically leads to sleep fragmentation and causes dysregulation of genes believed to play a role in sleep homeostasis. Our data also suggest that AMPKß loss may contribute to the increased risk of developing mental disorders and sleep disturbances associated with the human 1q21.1 deletion. Sleep is regulated by circadian and homeostatic processes. While the circadian clock is well studied, the molecular mechanism underlying homeostasis remains largely unknown. Our data suggest that AMPK is required for sleep maintenance and may be involved in the homeostatic process of sleep regulation.

**Author Summary:** The human 1q21.1 chromosomal deletion is associated with increased risk of schizophrenia. Because this deletion affects only a small number of genes, it provides a unique opportunity to identify the specific disease-causing gene(s) using animal models. Here, we report the use of a *Drosophila* model to identify the potential contribution of one gene affected by the 1q21.1 deletion – *PRKAB2* – to the pathology of the 1q21.1 deletion syndrome. *PRKAB2* encodes a subunit of the AMP-activated protein kinase (AMPK) complex, the main cellular energy sensor. We show that AMPK deficiency reduces lifespan and causes structural abnormalities in neuronal dendritic structures, a phenotype which has been linked to schizophrenia. Furthermore, cognitive impairment and altered sleep patterning are some of the most common symptoms of schizophrenia. Therefore, to understand the potential contribution of *PRKAB2* to the 1q21.1 syndrome, we tested whether AMPK alterations would cause defects in learning and sleep. Our study shows that lack of PRKAB2 and AMPK-complex activity, in the nervous system leads to reduced learning and to dramatic sleep disturbances. Thus, our data links a single 1q21.1-related gene with phenotypes that resemble common symptoms of schizophrenia, suggesting that this gene, *PRKAB2,* may contribute to the risk of developing schizophrenia.

## Introduction

Recent genome-wide association studies have identified copy-number variants (CNVs) associated with high risk of schizophrenia and other neuropsychiatric disorders [1–4]. Sleep disturbance is one of the most common problems in individuals with psychiatric disorders [5]. Although the 1q21.1 CNV deletion, spanning 10 genes over 1.35 million base pairs, has been demonstrated to genetically predispose carriers to sleep disturbances, intellectual disability, autism, and schizophrenia [4], the impact of individual genes in this interval on the development of these disorders is unclear. *PRKAB2,* encoding a ß-subunit of the AMP-activated protein kinase (AMPK) complex, is located on the 1q21.1 chromosome arm and is a promising candidate gene that may contribute to sleep disturbances and some of the behavioral effects observed in 1q21.1 patients. The AMPK complex is the main cellular energy sensor, functioning as a metabolic master switch that is required for maintenance of energy homeostasis [6]. When activated as a response to increased energy demands, through detecting increased intracellular AMP:ATP and ADP:ATP ratios, AMPK upregulates catabolic processes that generate adenosine triphosphate (ATP) while downregulating anabolic processes to maintain energy balance [7]. Interestingly, energy metabolism dysfunction is among the most consistent features observed in psychiatric disorders and has been shown to include pathways for ATP production [8]. The evolutionarily conserved AMPK complex that regulates ATP comprises a catalytic (serine/threonine kinase) α-subunit, a scaffolding ß-subunit, and a regulatory γ-subunit [9, 10]. Unlike mammalian genomes, which encode multiple isoforms of each AMPK subunit (α1, α2; ß1, ß2; γ1, γ2) that are, at least in part, functionally redundant, the fruit fly *Drosophila melanogaster* has only one gene encoding each subunit, encoded by *AMPKα, alicorn (alc),* and *SNF4Aγ,* respectively, making this organism an ideal model in which to study AMPK function [11].

While the function of AMPK in whole-body metabolism and cellular energy sensing is well-known, its role in neuronal maintenance, and the manner in which nervous-system-specific effects of AMPK manifest in global phenotypes, are poorly defined. Neurons in particular may be vulnerable to disturbances in AMPK function, as they have high metabolic demands and are ineffective at storing energy [12] – the human nervous system accounts for at least 20% of the body’s energy consumption while only making up 2% of the body’s mass [13]. Studies in mice have shown that AMPK is activated in the brain upon introduction of stresses such as glucose deprivation, ischaemia, and hypoxia [14, 15]. Furthermore, loss of the *AMPKβ1* subunit in mice was shown to have strong effects on brain development and structure [16]. Studies using *Drosophila* have shown that the AMPK complex affects cellular homeostasis, longevity, and neurodegeneration [17–19]. In *Drosophila,* upregulation of AMPKα in the nervous system has been linked to a slowing of systemic aging and thus to prolonged lifespan [20], which indicates that AMPK function in the nervous system has a broad, systemic impact. Furthermore, *alc,* encoding the single *Drosophila* AMPKβ homolog, has been shown to have a neuro-protective role in the retina [19]. However, a more generic role for the AMPK complex in the nervous system, and any downstream impacts on behavior, remain largely unknown.

To investigate the potential contribution of *PRKAB2* to the 1q21.1 CNV syndrome and the role of AMPK signaling in neuronal maintenance and sleep regulation, we examined the function of its *Drosophila* ortholog, Alc/AMPKβ, in the nervous system. Here, we show that *alc* and AMPK signaling are required in the nervous system to maintain behaviors such as sleep and learning. Specifically, we observed reduced learning in animals with neuronal knockdown of *alc.* Furthermore, we find that neuronal knockdown of *alc* and loss of AMPK function cause severe sleep fragmentation, and animals with reduced nervous-system *alc* expression are vulnerable to stress induced by sleep deprivation and show no sleep rebound. We suggest that *PRKAB2* is a promising candidate gene that may contribute to behavioral disturbances seen in 1q21.1 patients, such as sleep disturbances and cognitive deficits related to learning.

## Results

### Reduced *alc* in the nervous system shortens overall lifespan and increases sensitivity to starvation

Although AMPK activity has been linked to lifespan, the effect on longevity of reducing AMPK signaling specifically in the brain has not been reported. To determine whether neuronal reduction of AMPK signaling impacts overall physiology and health in *Drosophila,* we used the panneuronal *elav-GAL4* (*elav>*) driver to express either RNAi against *alc* (*alc-RNAi*; two independent constructs, in conjunction with Dicer-2) or overexpression of a kinase-dead dominant-negative form of the AMPKα subunit (*AMPKα-DN*)[17], and we monitored survival for 89 days. Both manipulations resulted in reduced survival compared to the driver (*elav>Dcr-2/+*) and corresponding RNAi (*alc-RNAi/+*) and *AMPKα-DN/+* controls (Fig 1A). To quantify lifespan, we fit the survival curves for individual experimental vials to a Weibull distribution to obtain a Weibull scale parameter that is analogous to the median survival but takes into account the increasing risk of mortality in aging systems (S1 Fig). Using this measurement, we found a significant reduction of lifespan in animals with nervous-system-specific knockdown of *alc* or overexpression of *AMPKα-DN,* conditions that reduce neuronal AMPK signaling (Fig 1B). This shows that effects of AMPK on lifespan can be attributed to its function in the nervous system.

**Fig 1.**
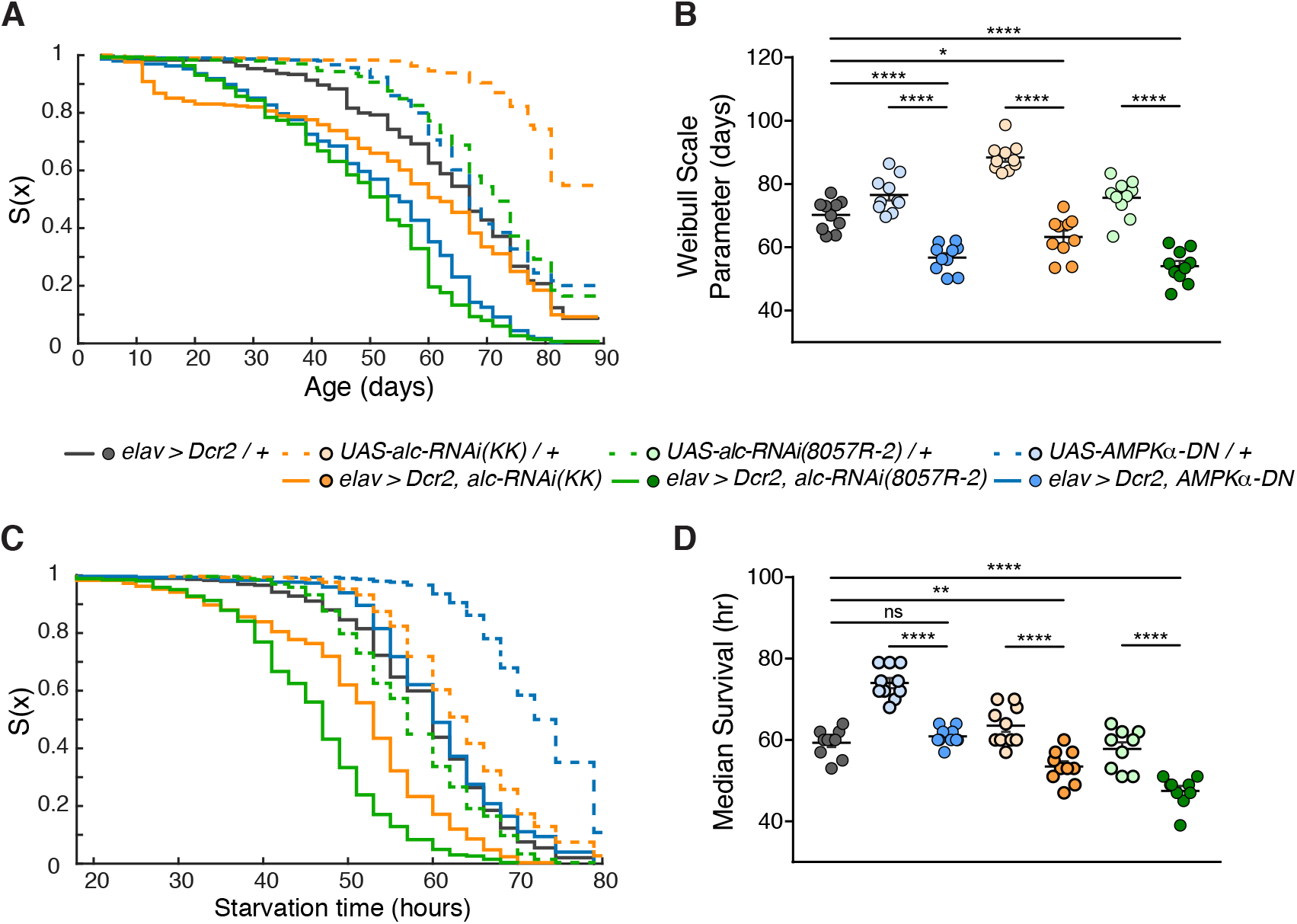
Loss of AMPK in the nervous system reduces overall lifespan. (A) Survival graph showing overall survival for an 89-day period, for male flies raised and kept on standard food. Results obtained from 10 replicates per genotype, each containing 30 flies. (B) Quantification of overall survival. Knockdown of *alc* or overexpression of *AMPKα-DN* in the nervous system reduces overall longevity. Survival analyzed using Kaplan-Meier nonparametric method. Survival quantified by fitting each replicate by a Weibull distribution function, right-censoring animals alive at end of experiment. (C) Survival graph showing overall survival for an 80-hour period, for 3-5-day-old male flies raised on normal food and starved on 2% agar alone. Results were obtained from 9-10 replicates per genotype, each containing 30 flies (n > 270). (D) Quantification of survival following starvation. Pan-neuronal knockdown of *alc* increases sensitivity to starvation. Error bars indicate SEM. Kruskal-Wallis test with Dunn’s post-hoc testing was used to determine statistical significance: *p<0.05, **p<0.01, ***p<0.001, ****p<0.0001, versus the control.

Several studies have reported an increased sensitivity to starvation in addition to decreased longevity in animals with reduced AMPK signaling [17, 21]. This effect has been speculated to involve metabolic pathways in adipose tissue, muscle, and intestine, since AMPK plays an important role in maintaining energy homeostasis especially under conditions of metabolic stress [22]. To investigate whether this reduced survival under starvation stress can be attributed to effects of AMPK in the nervous system, we determined the survival of animals with reduced neuronal AMPK signaling (knockdown of *alc* or overexpression of *AMPKα-DN*) under starvation. As we did in lifespan measurements, we observed reduced starvation survival with pan-neuronal *alc-RNAi*-mediated knockdown compared to controls (Fig 1C and 1D). Although animals overexpressing *AMPKα-DN* in the nervous system exhibited reduced survival compared to the *AMPKα-DN/+* control, we observed no difference compared to the *elav>Dcr-2/+* driver control. Together our results show that AMPK-mediated neuronal homeostasis plays a significant role in promoting organismal longevity and resistance to starvation. For the subsequent experiments, we chose to use the *UAS-alc-RNAi (KK*) line, which reduces expression of *alc* by ~85% in heads (S2A Fig). To further check the specificity of this RNAi towards *alc,* and knockdown of the AMPK complex in general, we examined phosphorylated AMPK*α* (pAMPK*α*) subunit levels, a readout of AMPK-complex activation. We found that neuronal knockdown of *alc* (AMPKß) caused a ~60% reduction in pAMPKα levels in the head, confirming that *alc* knockdown using this RNAi line specifically reduces AMPK activity (S2B Fig).

### *alc* is required for maintenance of dendritic arbors

The reduced lifespan observed in animals with neuronal loss of AMPK signaling suggested that lack of AMPK activity may disrupt energy-homeostatic processes and lead to impaired neuronal function and maintenance. Since changes in neuronal dendritic branching and morphology are a hallmark of neuropsychiatric disorders and have been suggested to alter the function of neuronal circuitry [23–26], we chose to analyze the effect of *alc* manipulations on dendritic development and maintenance. In *Drosophila,* the class-IV dendritic arborization (da) neurons are peripheral sensory neurons that develop highly stereotyped dendritic processes, making them an ideal model for studying changes in neuronal morphology[27]. Disruption of *alc* has been shown to cause progressive and activity-dependent degeneration of photoreceptor neurons[19], but its impact on dendritic arborization has not been explored. To determine whether *alc* has a role in dendrite development or maintenance, we reduced expression of *alc* using the da-neuron-specific *pickpocket (ppk)-GAL4* (*ppk>*) driver line while simultaneously expressing *GFP* to visualize neuronal structures. To obtain higher-quality data, we imaged individual class-IV da neurons in both large feeding (~100 hours after egg lay) and wandering (~120 hours after egg lay) 3^rd^-instar larvae. RNAi-mediated knockdown caused a dramatic reduction of dendritic arborization in feeding larvae, reducing the total dendrite length by 25% (Fig 2A and 2B) and branching order by 14.5%, limiting dendrite structure to mainly primary and secondary branches (Fig 2C). Knockdown of *alc* also led to 35% fewer dendritic branches yet did not alter the total area spanned by individual neurons, compared to controls (Fig 2D and 2E). The severity of this dendritic phenotype was progressive, appearing more severe in older (wandering) animals, with a 44% reduction in total dendrite length (Fig 2B), 38% reduction in branch order (Fig 2C), and 68% reduction in branch number (Fig 2E). This observation suggests that AMPK is required for the maintenance of dendritic structures, rather than for their formation. Furthermore, we observed a “beaded” dendritic morphology as well as thinned and fragmented primary dendritic branches, which are hallmarks of dendritic degeneration. Our AMPKß observations in da neurons are further supported by previous observations of similar dendrite-morphological abnormalities associated with loss of AMPKα and AMPKγ subunit activity in these cells [17]. Together these observations suggest that AMPK activity is required to prevent energy deficiency in neurons and that this process is necessary for the maintenance of dendritic arbors. Since proper dendritic structure is required for a functional nervous system, we then asked whether AMPK signaling, and specifically *alc*, may be required for proper expression of behaviours relevant to the development of neurological disorders.

**Fig 2.**
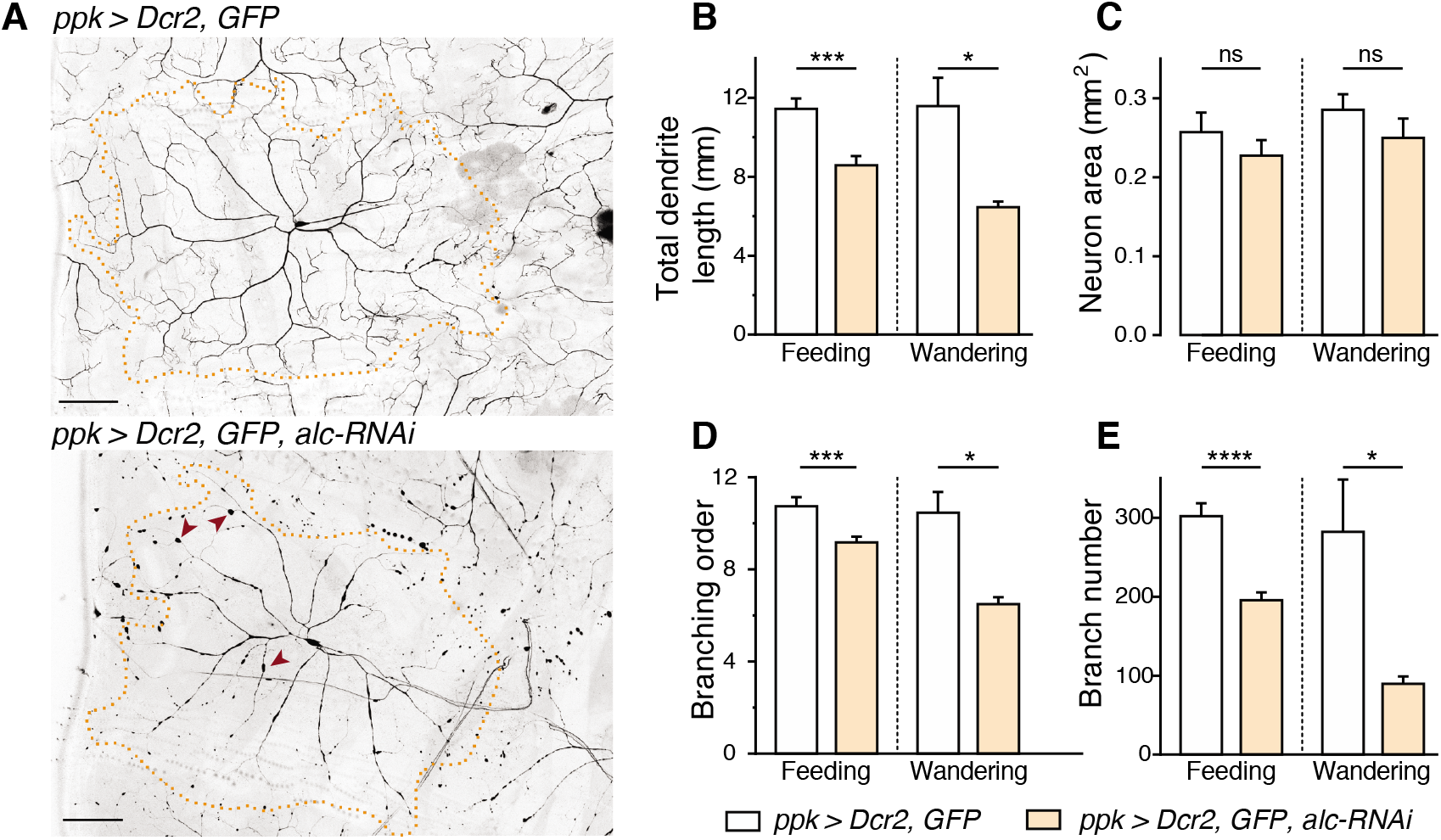
*alc/AMPKβ* is required for maintenance of dendritic branching in Class-IV sensory neurons. (A) Representative images of Class-IV *pickpocket* (*ppk)>GFP*-expressing neurons from abdominal segment A2 in control (top) and *alc-RNAi* knockdown third-instar larvae. Individual neurons are outlined. Arrowheads point to beaded morphology of dendrites characteristic of reduced *alc.* (B) Quantification of total dendrite length (mm) in both feeding (n = 10) and wandering (n = 3) third-instar larvae shows a significant decrease in total dendrite length when *alc* is knocked down in class-IV neurons. (C-E) Quantification shows no significant change in neuronal area (mm^2^) (D); however, knockdown of *alc* decreases both dendritic branching order (C) and branch number (E). Quantification performed using the TREES Matlab toolbox. Error bars indicate SEM. Unpaired T-test was used to determine statistical significance: *p<0.05, **p<0.01, ***p<0.001, ****p<0.0001, versus the control.

### *alc* is required for learning

Intellectual disability and general learning disabilities are prevalent in patients carrying the 1q21.1 CNV [4, 28]. Since *alc* is required for proper dendritic arborization of neurons, we hypothesized that *alc* might also be important in the maintenance of behaviors such as learning. In *Drosophila,* one simple behavioral assay that tests learning is courtship conditioning [29]. In this experiment, naïve male flies are presented with an unreceptive mated female and soon associate her mating-associated olfactory and gustatory cues with the courtship-rejection behaviors she expresses, leading the males to suppress futile courtship behavior towards other mated females in the future. To determine whether *alc* is required for learning following courtship conditioning, we conducted a courtship-conditioning assay with adult (4-7 day old) flies. Following a 1-hour training session in which naïve males were presented with a mated (unreceptive) female, individual naïve or trained males were introduced to a mated female. Animal pairs were video-recorded using a custom recording setup in custom chambers (S3 Fig), and stereotypical courtship behaviors were manually scored. Courtship indices (CIs) were calculated as the fraction of time each male spent courting within the 10-minute recording period. Males from control genotypes (*elav>Dcr-2/+* and *alc-RNAi/+*) showed typical levels of courtship suppression following training, compared to naïve flies exposed to a sham training session (Fig 3A). In sharp contrast, there was no significant difference in CI between naïve and trained *elav>Dcr-2,alc-RNAi* flies with reduced expression of *alc* in the nervous system, indicating that pan-neuronal knockdown of *alc* results in inability to suppress courtship behavior. The percentage reduction in CI of naïve versus trained flies can be reflected in the learning index (LI), which is essentially zero in animals with reduced neuronal expression of *alc* (Fig 3B). These data suggest that neuronal expression of *alc* is required for learning during courtship conditioning.

**Fig 3.**
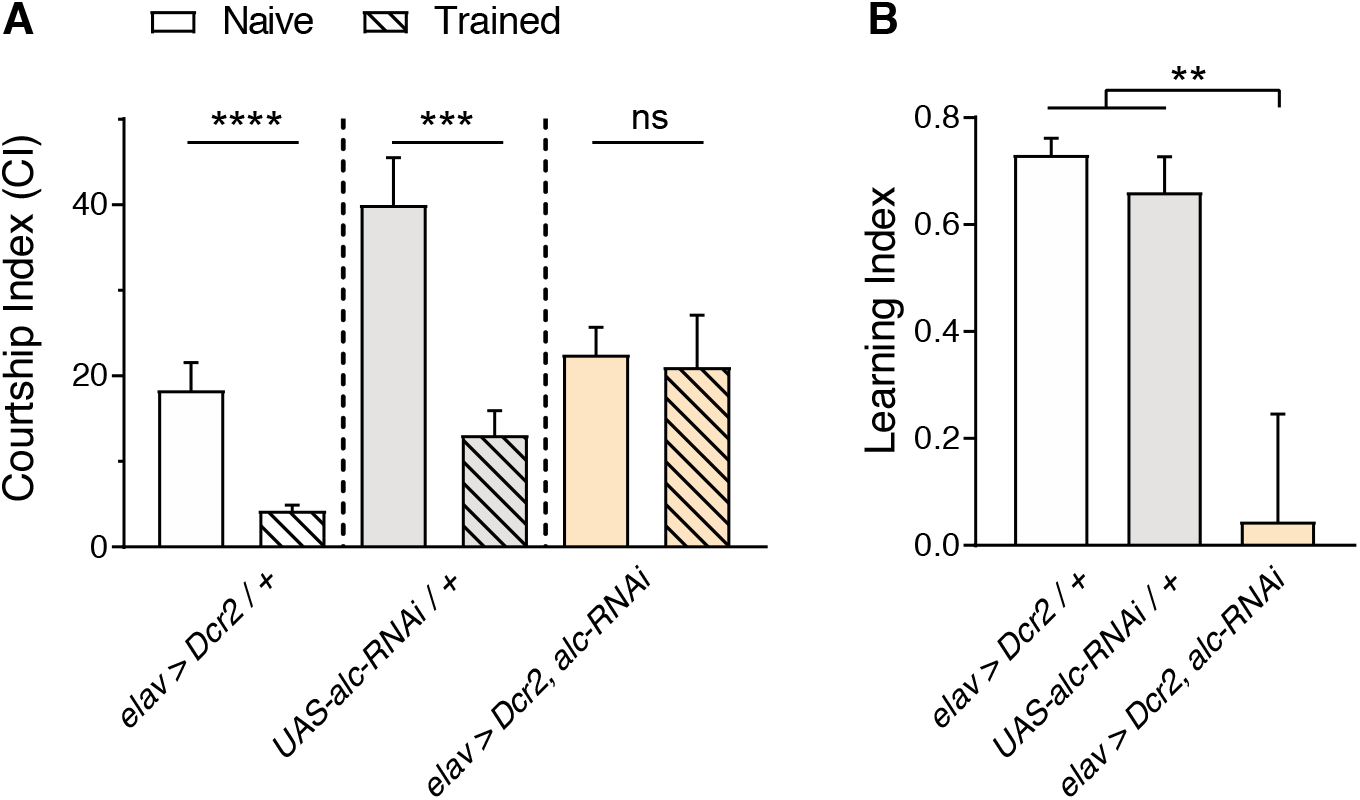
Knockdown of *alc/AMPKβ* in the nervous system impairs courtship-conditioning learning. (A) A courtship-conditioning assay was used to determine whether *alc* is required for courtship learning. Courtship index (percentage of time spent courting) decreased after training (with a mated female) in control animals (77% decrease for *elav>Dcr-2/+* and 67% decrease for *UAS-alc-RNAi/+*), indicating learning in both genotypes. There was no significant difference between the performance of naïve and trained animals lacking *alc* pan-neuronally. Mann Whitney test was used to determine statistical significance: **p<0.01, ***p<0.001, ****p<0.0001, versus the control. Results were obtained from 11 experimental repeats for each genotype, using 3 animals trials each (n = 33 animals). (B) Calculated learning index for *alc* knockdown and control animals. Error bars indicate SEM. Significance was determined using a one-way ANOVA with Tukey’s post-hoc comparison: **p<0.01 versus the control.

### Loss of *alc* in the nervous system disrupts sleep architecture

Patients with the 1q21.1 microdeletion also suffer from non-intellectual behavioral deficits, a common manifestation being disrupted sleep [30]. It has been suggested that sleep is essential for the brain to replenish energy sources (ATP) that were depleted during wakefulness [31]. As one of the major roles of the AMPK complex is to upregulate processes that generate ATP, we asked whether knockdown of *alc* in the nervous system would impair sleep consolidation. We hypothesized that if sleep drive is associated with the depletion of energy stores during metabolically demanding wakefulness, thereby stimulating AMPK-dependent signaling, levels of activated AMPK would fall during sleep as ATP levels are restored. Under this scenario, a decrease in AMPK activity would be associated with decreased sleep drive, increased wakefulness, and reduced consolidation of sleep episodes. Sleep in *Drosophila* can be measured using the *Drosophila* Activity Monitor (DAM) system (TriKinetics), an automated beam-crossing locomotion assay that is the widely accepted standard in *Drosophila* sleep studies. It is well established in *Drosophila* that periods of locomotor quiescence of longer than 5 minutes indicate a sleep-like state sharing the hallmarks of mammalian sleep, such as increased arousal threshold [32, 33] To determine whether loss of *alc* impairs sleep, we expressed *alc-RNAi* in the nervous system and monitored sleep using the DAM system in a light/dark illumination-controlled incubator. Animals with reduced *alc* expression in the nervous system exhibited significantly reduced overall sleep compared to control genotypes, during both day and night phases. This is also reflected in an overall increase in activity compared to controls (Fig 4A–4C). Thus reduced “sleep,” defined as it is as long-term immobility, can be indicative of a hyperactivity phenotype. To clarify this, we examined the properties of activity bouts between periods of sleep. As *Drosophila* spends circadian light transition events in an elevated state of arousal, we excluded these periods from our analyses. Activity during wakeful periods (number of beam crosses per minute awake) was significantly increased during the light phase, but not during the dark phase (Fig 4D). Flies also demonstrated a significant increase in the duration of wakeful periods during light (compared to both controls) and dark (compared to *UAS-alc-RNAi/+* control only) (Fig 4E). Interestingly, when we analyzed the periods around light-transition events – periods of elevated locomotive activity – we did not observe increased level or duration of activity for *elav>Dcr-2,alc-RNAi* flies compared to controls (S4 Fig), which indicates that the loss of *alc* causes a specific sleep phenotype that is not a consequence of overall hyperactivity. Together these data suggest that animals with reduced expression of *alc*, and thus reduced AMPK activity, in the nervous system have an increased drive for wakefulness and reduced sleep drive.

**Fig 4.**
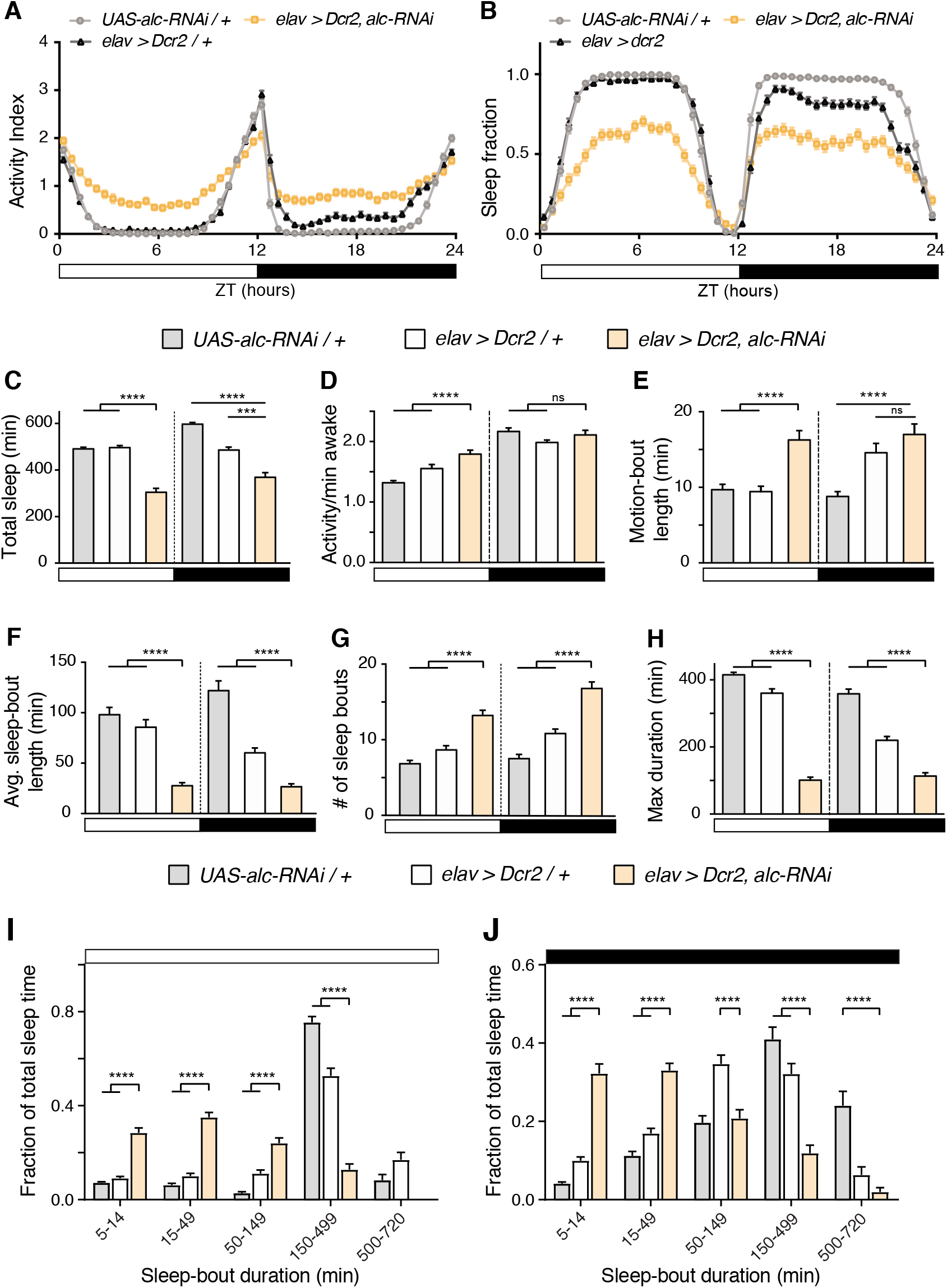
Knockdown of *alc/AMPKβ* in the nervous system disrupts sleep. (A-B) Activity (A) and sleep (B) profiles over a 24-hour period for control genotypes *elav>Dcr-2* (n = 128) and *UAS-alc-RNAi/+* (n = 126) versus *elav>Dcr-2, alc-RNAi* (n = 126). All data obtained from second 24-hour cycle to allow for acclimatization. Activity and sleep are shown in bins of 30 minutes. White and black bars represent ZT time, 12 hours light and 12 hours dark, respectively. (C) Total sleep (min) in flies with pan-neuronal *alc* knockdown compared to controls. Total sleep is significantly reduced when *alc* is knocked down in the nervous system. (D) Comparison of activity during outside sleep periods between *alc* knockdown animals and controls shows that pan-neuronal knockdown of *alc* increases activity during daytime, but not nighttime. (E) Motion-bout length (min) is significantly increased during the day in *alc* knockdown flies compared to controls. (F) Average sleep-bout length (min) is significantly reduced during both day- and night-time in *alc* knockdown flies. (G) The number of sleep bouts per day and night increases when *alc* is knocked down in the nervous system. (H) Duration of the longest sleep bout (min) is significantly shorter in *alc* knockdown animals than in controls. (I, J) Distribution of length of sleep bouts for control genotypes *elav>Dcr-2/+* and *UAS-alc-RNAi/+* versus *elav>Dcr-2, alc-RNAi* during the day (I) and during the night (J). Flies in which *alc* has been knocked down pan-neuronally only generate a small proportion of sleep bouts of >150 min, whereas the majority of sleep bouts in controls are of >150 min. Error bars indicate SEM. Kruskal-Wallis test with Dunn’s post-hoc testing was used to determine statistical significance: ***p<0.001, ****p<0.0001, versus the control.

To further assess the sleep properties of animals with neuronal loss of *alc* function, we investigated sleep-bout duration and number. Animals with reduced *alc* expression in the nervous system showed an increase in sleep bout-number and a concomitant significant reduction in average sleep-bout length, indicating highly fragmented sleep (Fig 4F and 4G). The average longest single sleep bout during both day and night was also greatly reduced compared to control genotypes (Fig 4H). To more closely examine these sleep changes, we analyzed the distribution of sleep episodes by binning them according to duration and calculating the time spent in each bin. Whereas control flies consolidated most of their daytime and nighttime sleep into long sleep episodes of 150-499 minutes, animals with neuronal loss of *alc* demonstrated a shifted sleep structure with significantly more time spent in short sleep bouts (binned into 5-14 and 15-50 minutes) (Fig 4I and 4J). In the aggregate, these data indicate that *alc* is necessary for proper sleep architecture and consolidation in *Drosophila*.

### Loss of AMPK signaling causes progressively worsening sleep fragmentation

To confirm the reduced-sleep phenotype observed for pan-neuronal knockdown of *alc,* we used a second, independent *alc-RNAi* line (*8057-R2*) line and determined average sleep-bout duration and number during both day- and night-time. As seen with the *alc-RNAi(KK*) line, sleep-bout duration was significantly decreased in length during both circadian phases, and sleep-bout numbers were greatly increased over the 24-hour period, reflecting increased sleep fragmentation (Fig 5A and 5B). To examine whether this phenotype was specifically due to altered AMPK signaling in the nervous system, we overexpressed *AMPKα-DN* to block AMPK-complex activity via inhibition of the alpha subunit. Surprisingly, these animals initially exhibited no significant reduction in average sleep-bout duration or number. However, over longer observation periods, sleep phenotypes arose that were similar to those displayed by alc-knockdown animals – average sleep-bout duration significantly decreased over time, while the number of sleep bouts significantly increased, during both day and night (Fig 5C and 5D). This progressive deterioration in sleep consolidation over time was specific to reduced AMPK signaling, since no significant changes in sleep-bout architecture were observed with control genotypes aside from a slightly *increased* daytime sleep-bout duration in one control line. Thus, the loss of AMPK signaling in the nervous system causes sleep fragmentation that worsens over time. This progressive deterioration is reminiscent of the morphological phenotypes observed in larval class IV da neurons. It is possible that a similar morphological deterioration of sleep-regulatory neurons might underlie these defects.

**Fig 5.**
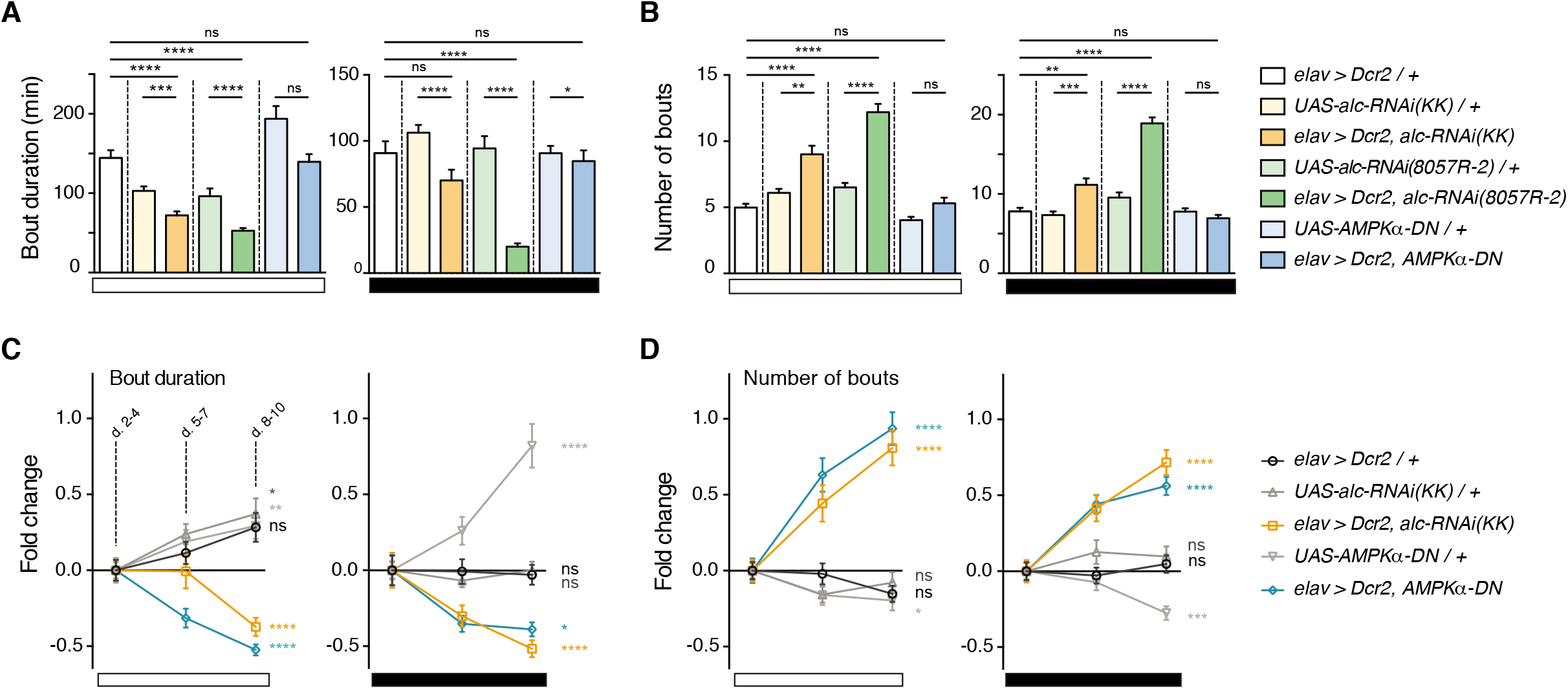
Neuronal loss of AMPK activity causes progressive sleep fragmentation. (A-B) Quantification of average bout durations (A) and average bout numbers (B) during day and night for *elav>Dcr-2/+* (n = 32), *UAS-alc-RNAi(KK)/+* (n = 32), *elav>Dcr-2, alc-RNAi(KK*) (n = 31), *UAS-alc-RNAi(8057-R2)/+* (n = 32), *elav>Dcr-2, alc-RNAi(8057-R2*) (n = 32), *UAS-AMPKα-DN/+* (n = 32), *elav>Dcr-2, AMPKα-DN* (n = 31). Sleep was monitored over 10 days, and data from days 2 to 4 was used. Bout durations are significantly reduced and bout numbers significantly increased when *alc* is knocked down in the nervous system compared to control genotypes, for both RNAi lines. There is no initial difference in either parameter when dominant-negative *AMPKα (AMPKα-DN*) is overexpressed in the nervous system. (C-D) Graphical representation of the fold change in bout duration (C) and bout number (D) over time (3-day binned) during both day and night, indicated by white and black bars respectively. Shown are data for *elav>Dcr-2, alc-RNAi(KK*) and *elav>Dcr-2, AMPKα-DN* compared to control genotypes. Bout duration significantly decreases over time in both *alc* knockdown and *AMPKα-DN* overexpression flies. Bout number significantly increases over time for both genotypes. Significance between days 8-10 and days 2-4 is shown. Error bars indicate SEM. Significance was determined using a Kruskal-Wallis test with Dunn’s post-hoc testing: *p<0.05, **p<0.01, ***p<0.001, ****p<0.0001, versus the control.

### Animals defective in AMPK signaling are in a state of sleep deprivation

Since defective neuronal AMPK signaling due to tissue-specific *alc* knockdown resulted in fragmented and reduced sleep, we suspected that these animals might therefore exhibit altered expression of sleep-stress-response genes. To assess this possibility, we performed RNA sequencing on adult head samples of animals with neuronal knockdown of *alc* versus controls. Amylase has been identified as a biomarker for sleep drive in *Drosophila*, and another, previously uncharacterized gene, *heimdall,* has recently been linked to animals’ response to sleep deprivation [34, 35]. Interestingly, both of these genes are significantly upregulated in the heads of *alc*-knockdown flies – *heimdall* exhibited the greatest transcriptional up-regulation of any gene, with a roughly 180-fold increase compared to control heads (Table 1 and S1 Table). Thus, alc-knockdown animals show a genetic response consistent with disrupted sleep.

**Table 1.** RNA-seq data for top differentially expressed genes in adult male fly heads when *alc* is knocked down in the nervous system. Fold change in transcript levels for *elav>Dcr2, alc-RNAi(KK*) compared to the *elav>Dcr2/+* controls.

| Annotation | Gene Name | Fold Change | P-Value |
| --- | --- | --- | --- |
| CG4500 | heimdall | 179.53 | 8.32E-257 |
| Hsp70Bb | Heat-shock-protein-70Bb | 45.15 | 0 |
| CG30059 | - | 34.17 | 4.03E-32 |
| CR44994 | - | 33.82 | 2.66E-08 |
| Mal-A8 | Maltase A8 | 30.60 | 5.81E-12 |
| CG34040 | - | 22.44 | 6.27E-06 |
| Mal-A7 | Maltase A7 | 20.80 | 4.81E-12 |
| Cyp6d2 | Cyp6d2 | 15.82 | 7.14E-45 |
| lectin-24A | lectin-24A | 15.52 | 3.16E-07 |
| squ | squash | 11.89 | 8.54E-22 |
| Amy-p | Amylase proximal | 8.71 | 7.56E-09 |
| alc | alicorn | -2.50 | 0 |
| Obp56a | Odorant-binding protein 56a | -6.55 | 5.90E-161 |
| CG32437 | - | -6.90 | 4.60E-07 |
| CG8563 | - | -6.94 | 7.89E-15 |
| Obp99d | Odorant-binding protein 99d | -14.39 | 2.68E-36 |
| CG43291 | - | -18.69 | 9.75E-24 |
| CG12971 | - | -35.38 | 1.26E-30 |
| CG17242 | - | -40.90 | 2.58E-05 |
| CR45600 | - | -62.45 | 0 |

### *alc* is required for initiation and maintenance of “recovery sleep” following sleep deprivation

A major hallmark of sleep is homeostatic regulation, which is required for the recovery of lost sleep after sleep deprivation (SD). To investigate whether AMPK is important for the homeostatic process of sleep regulation, we tested whether animals expressing neuronal *alc* RNAi were defective in recovery sleep following deprivation. To assess recovery sleep following SD in *alc* knockdown animals, we exposed *elav>Dcr-2/+* control animals and *elav>Dcr-2,alc-RNAi* animals to mechanical SD, using a vortex mounting plate (Trikinetics) that regularly jostled the animals. Sleep deprivation was initiated during the second day of recording, for the 6 hours of the latter half of the dark phase immediately preceding “lights-on.” This resulted in similar levels of sleep loss in *alc* knockdown and control animals (Fig 6A, and 6B). However, whereas driver controls showed substantial subsequent sleep rebound following SD, *elav>Dcr-2,alc-RNAi* animals did not show any such post-SD rebound (Fig 6B and 6C). Following relief from SD, control flies rapidly fell into extended sleep and entered their longest consolidated sleep episode faster (Fig 6D), with an increase in average bout duration initiated during the first two hours following SD (Fig 6E). In contrast, animals lacking *alc* in the nervous system took significantly longer to reach a maximally consolidated sleep bout and did not increase sleep bout duration following SD (Fig 6D and 6E). This indicates that animals with impaired neuronal AMPK signaling not only fail to initiate rebound sleep, but also seem to be generally impaired in their sleep following SD. To further examine this deprivation-dependent shift in sleep architecture, we analyzed the distribution of sleep bouts during day and night following the deprivation period. Directly following sleep deprivation, control animals spent more time in prolonged sleep episodes (500-720 min) compared to baseline periods, and returned to baseline sleep-length distribution in the following dark phase (Fig 6F). In contrast, *elav>Dcr-2,alc-RNAi* animals displayed more-fragmented sleep immediately following deprivation, indicative of an impaired sleep drive (Fig 6G). Surprisingly, this fragmentation persisted into the following dark phase, indicating that SD of flies with altered AMPK signaling has longer-term effects on sleep architecture (Fig 6G).

**Fig 9.**
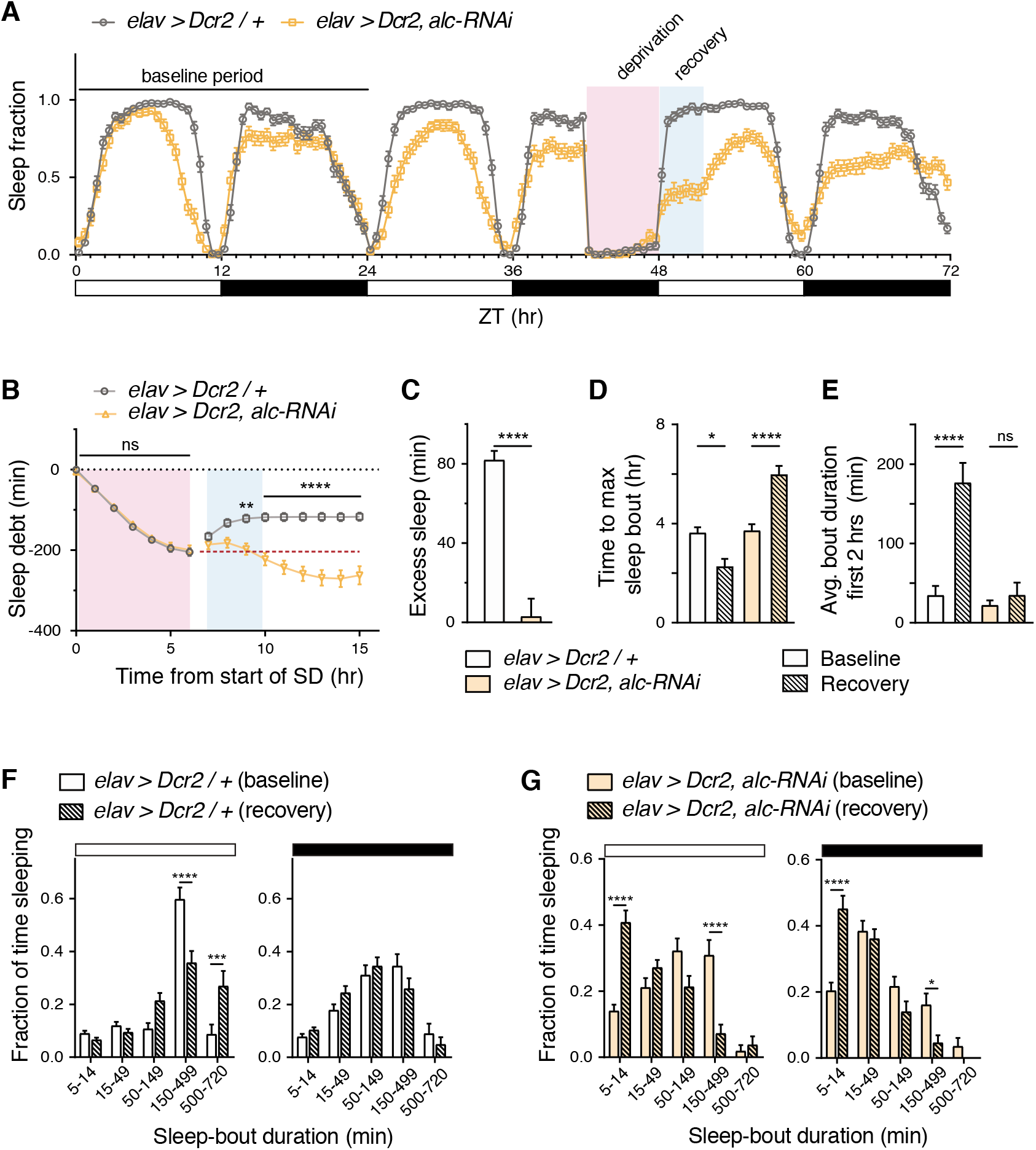
Alc/AMPKβ is required in the nervous system for recovery sleep following sleep deprivation. (A) Sleep profile for control genotype *elav>Dcr2/+* (n = 54) versus *elav>Dcr2, alc-RNAi* (n = 52) for a 72-hour period, showing sleep profiles for baseline, sleep deprivation, and recovery days. White and black bars represent ZT time, 12 hours light and 12 hours dark, respectively. A 6-hour sleep-deprivation period is illustrated using pink shading, typical recovery period illustrated with blue shading. (B) Quantification of sleep debt (min) showed that control flies recovered lost sleep in the first 3 hours following sleep deprivation. Significance determined by 2-way ANOVA with Sidak’s multiple comparisons test between control and *alc-RNAi* animals shown. Dashed line shows maximum sleep debt. (C) Control animals showed a significant excess of sleep (80 minutes) compared to the same time period of the baseline day while *alc-RNAi* animals did not. Pan-neuronal knock down of *alc* completely eliminates recovery sleep. (D) Time to maximum sleep bout (hours) is significantly reduced in control flies following sleep deprivation, whereas it is significantly increased in flies lacking *alc* in the nervous system. (E) Average bout duration in the first two hours of light phase was significantly increased during recovery in control flies, but not in *alc* knockdown animals. (F-G) Distribution of length of sleep bouts for control genotype *elav>Dcr2/+* (F) versus *elav>Dcr2, alc-RNAi* (G) during the day and during the night, for baseline and recovery days. Unlike control flies, which showed an increase in long sleep bouts (>150 minutes) during the day, *alc* knockdown animals showed a decrease in long sleep bouts during both the day and the night, following sleep deprivation. Error bars indicate SEM. For C-E, Mann Whitney test was used to determine statistical significance. For F and G, a 2-way ANOVA with Tukey’s post-hoc comparison was used: *p<0.05, **p<0.01, ***p<0.001, ****p<0.0001, versus the control.

To determine whether the apparent detrimental effect of sleep deprivation impacts survival, we monitored sleep-deprived animals for 24 hours following sleep deprivation. We observed increased mortality of *elav>Dcr-2,alc-RNAi* animals following sleep deprivation compared to both un-deprived age-matched *elav>Dcr-2,alc-RNAi* animals and SD and non-SD driver controls (S5A Fig). To control for a possible elevated sensitivity to mechanical stress, we performed a mechanical stress assay in which flies were exposed to prolonged repeated mechanical perturbation and stress-induced mortality was assessed [36]. Knockdown of *alc* in the nervous system did not result in a higher incidence of lethality over this period (S5B Fig), which confirms that the observed lethality following sleep deprivation and the corresponding deterioration of sleep architecture is the result of sleep loss rather than physical weakness of the animals. Together, these data indicate that *alc* knockdown animals are incapable of consolidating rebound sleep and are highly sensitive to sleep deprivation, with long-lasting detrimental effects following SD.

Our findings indicate that AMPK activity is required for rebound sleep following deprivation, indicating that the AMPK complex is involved in the homeostatic regulation of sleep. In *Drosophila*, a brain structure known as the central complex plays a key role in sleep homeostasis [37–40], so we therefore investigated whether *alc* is required in this structure for regulation of sleep. First, to examine the distribution of Alc within the brain, we generated a fluorescent Venus-tagged form of the endogenous Alc protein using CRISPR. We observed Alc::Venus signal in the brain with specific labeling of the central complex, which was diminished in *alc-RNAi* animals (S6 Fig), consistent with a function of Alc in homeostatic features of sleep regulation. We next investigated whether loss of *alc* specifically in the neurons of the central complex affects sleep. We used a panel of three drivers expressing GAL4 in the central complex, including *R24C10-GAL4,* which was recently shown to drive expression in neurons that play important roles in sleep homeostasis [41, 42]. Driving *alc-RNAi* using these drivers lead to a fragmentation of sleep with more bouts of shorter duration, similar to that observed with pan-neuronal knockdown (Fig 7A and 7B). This supports a role for Alc in neural circuitry that is involved in homeostatic regulation of sleep. According to this view, AMPK activity would increase over periods of wakefulness, when higher neuronal energy demands deplete cellular energy stores. Consistent with this, we observed upregulation of pAMPKα levels in heads of control animals after 6 hours of SD, whereas this signal did not increase in animals with reduced neuronal *alc* expression (Fig 7C). Elevated pAMPKα levels were also observed in wild-type (Canton S) flies after SD, confirming AMPK-complex activation following periods of SD in wild-type animals.

**Fig 7.**
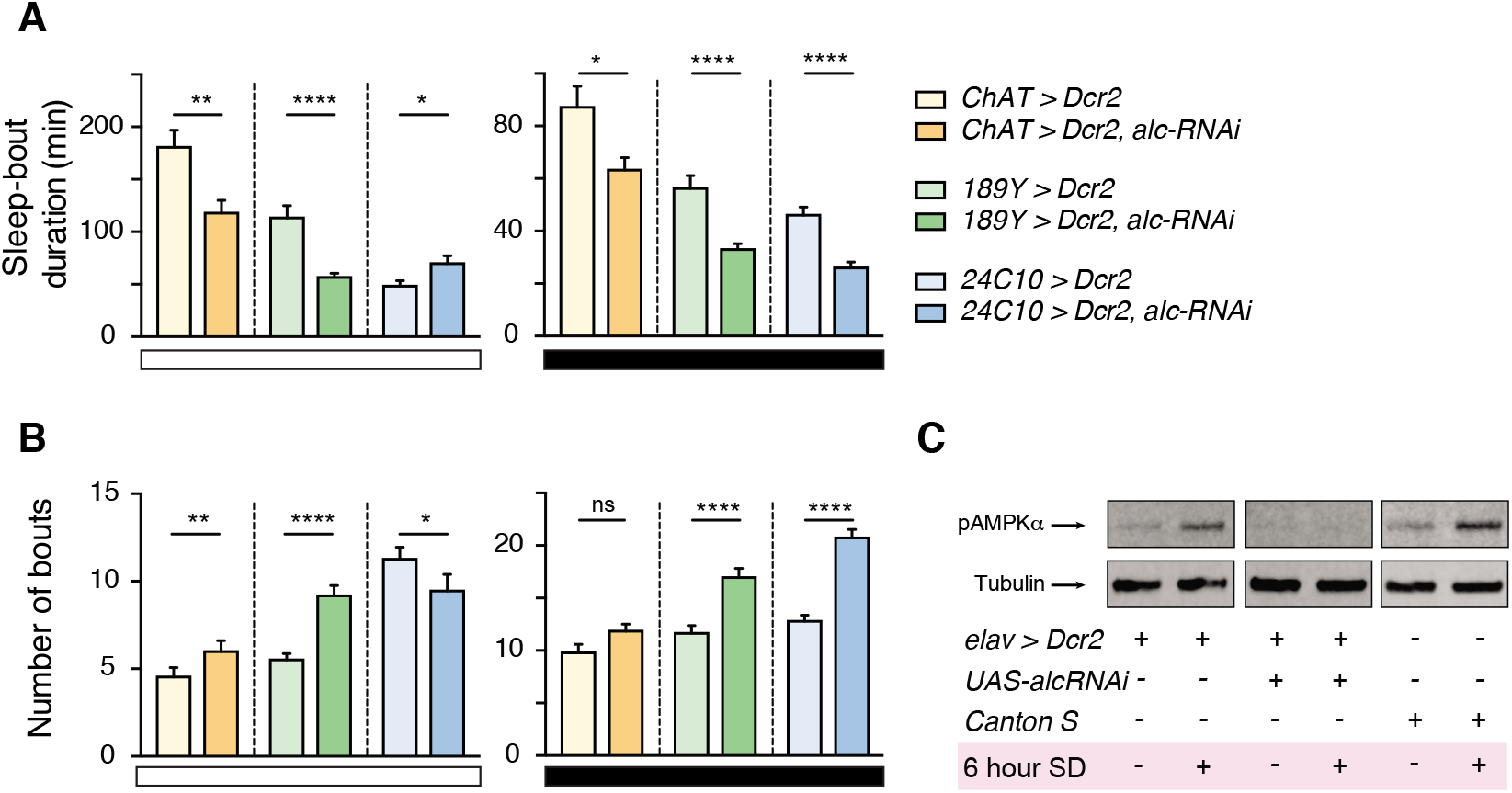
Loss of *alc/AMPKβ* in the central complex disrupts sleep. (A) Average sleep-bout length (min) was significantly reduced during both day- and night-time when *alc* was knocked down in the central complex using three previously described GAL4 lines, *ChAT-GAL4, 189Y-GAL4,* and *R24C10-GAL4,* with the exception of *R24C10>alc-RNAi* during the day. (B) The number of sleep bouts per day and night increased when *alc* was knocked down in the central complex. (C) Levels of phosphorylated AMPK*α* (pAMPK*α*) increased (normalized to alpha-Tubulin) immediately following 6 hours of sleep deprivation (SD) compared to same-time non-deprived animals for control animals (*elav>Dcr-2/+* and *Canton S*).

## Discussion

The shortening of sleep-bout duration and the associated increase in their number is indicative of a defect in sleep maintenance, in which sleep is both reduced and fragmented. Furthermore, perturbations of sleep structure are a hallmark of many neuropsychiatric and neurodegenerative disorders, and this type of perturbation is associated with the 1q21.1 deletion, which includes *PRKAB2/AMPKβ2* [1–4]. Here we show that Alc/AMPKβ and AMPK signaling are necessary for sleep maintenance, with reduced AMPK-complex activity resulting in fragmented sleep episodes. Our findings indicate that animals with disrupted neuronal AMPK signaling have an impaired ability to maintain the sleep state. The loss of post-deprivation rebound sleep in animals lacking neuronal AMPK activity further suggests a specific effect on sleep homeostasis. Because sleep serves beneficial biological functions, the persistent state of reduced sleep might also explain the shortened lifespan of flies with reduced AMPK signaling in the nervous system. The role of AMPK in stress resistance is well established [11], however, its role in the regulation of sleep and in the response to sleep deprivation has not been previously reported. The disruption of sleep parameters following deprivation, and the associated high level of mortality, indeed indicates a detrimental effect that may be associated with lack of rebound sleep. As increased neuronal activity under reduced AMPK signaling has been shown to result in degeneration of retinal photoreceptive neurons [19], it is possible that elevated activity during sleep deprivation, without the neuroprotective mechanisms of AMPK, leads to some degree of excitotoxicity. In any case, our *Drosophila* data suggest that *AMPKβ2* may underlie or contribute to the phenotypes observed in human 1q21.1 CNV syndromes. This provides a molecular mechanism for understanding the link between the deletion and increased risk of developing mental disorders associated with the 1q21.1 CNV, which may be important for development of new strategies aimed at treating both mental disorders and sleep disturbances.

### Neuronal AMPK activity affects lifespan and stress resistance

AMPK signaling has been shown to play a critical role in aging and lifespan determination [43]. Upregulation of AMPK signaling in *Drosophila* has been shown to extend lifespan by mediating effects specifically in the intestine and the brain [20]. Furthermore, tissue-specific RNAi-mediated knockdown of *AMPKα* in the fat body and muscle was shown to reduce lifespan [44, 45]. We find that AMPK signaling in the nervous system affects longevity and survival under starvation stress. This indicates that non-metabolic pathways may contribute to the impairment of survival upon loss of AMPK signaling. As non-cell-autonomous mechanisms have been implicated in the regulation of longevity upon enhanced AMPK signaling in the brain [20], it is possible that similar mechanisms are involved in the reduction of lifespan we observe. Interestingly, reduction of AMPK in neuroendocrine cells releasing the energy-mobilizing Adipokinetic Hormone resulted in a significant extension of life span under starvation [46]. An intriguing possibility is that the starvation sensitivity that we observed with nervous-system AMPK loss is mediated through effects on neuromodulatory or neurohormonal systems.

### AMPK is required for neuronal maintenance

The cellular mechanisms by which altered metabolism brings about neuropathology are not clear. It is well established that neuronal morphogenesis defects may result from mitochondrial dysfunction in various cell types [47, 48]. As mitochondria and AMPK are both involved in maintaining levels of ATP, defects in these systems may phenocopy each other. In fact, it has been shown that loss of AMPK activity enhances neurodegeneration in *Drosophila* models of mitochondrial abnormalities [49]. Normal neuronal activity and synaptic transmission involve molecular and cellular processes, such as maintenance of resting membrane potential and generation of action potentials, transporter activity, transmitter synthesis, and vesicle transport and dynamics, that are energy-intensive. Therefore, a disruption of restorative AMPK signaling that is required to replenish energy stores could result in a state of neuronal energy depletion, which could lead to progressive degeneration. Here we show that knockdown of *alc* results in progressive reduction of the dendritic arbors of class-IV da neurons, a phenotype consistent with reduced AMPK activity [17, 50]. Reducing AMPK signaling by knockdown of *alc/AMPKβ* specifically affects terminal dendrite branching, not the distance (reach) from the soma. Our observations are consistent with previous reports showing that altered AMPK signaling results in neurodegeneration [51, 52]. RNAi-mediated knockdown of the alpha or gamma subunits of the AMPK complex in class-IV da neurons in *Drosophila* larvae causes aberrant dendrite morphology, indicative of faulty neuronal development, neuronal damage, and degeneration. In sum, this suggests that neuronal loss of AMPK activity is associated with progressive neurodegeneration, originating with insufficient energy to maintain neuronal structures. These results are also consistent with the emerging role of AMPK in neurodegenerative diseases such as Alzheimer’s, Parkinson’s and Huntington’s [53].

### Requirement of Alc/AMPKβ for learning

The role of AMPK in mediating the synaptic plasticity that underlies learning and memory consolidation is unclear. One possibility to explain our observations would be that the neuronal connections necessary for associative learning in the mushroom body of the insect brain [54] have degenerated, thus resulting in a system incapable of forming, retaining, or recalling memories. Another possibility is that loss of AMPK may impair cAMP second-messenger signaling, which underlies the neuronal plasticity necessary for learning and memory [55, 56]. In mammalian adipocytes, AMPK has been shown to be activated by intracellular cAMP levels [57]; it is possible that a similar mechanism is present in neurons and is involved in mediating neuronal activity and plasticity. Interestingly, in mice, treatment with the AMPK agonist AICAR increases spatial memory in a Morris water maze [58, 59]. Our results suggest that Alc/AMPKβ and thus the AMPK complex are required for learning, although further experiments need to be conducted to evaluate the role of reduced AMPK signaling in the formation and maintenance of memory. Interestingly, it has been shown that sleep deprivation has a negative effect on learning and memory in several animal models [60–62]. A persistent state of sleep deprivation in animals deficient in AMPK signaling may explain their inability to learn to repress courtship upon rejection. Other learning paradigms need to be tested to determine if this is a general effect on the association center of the *Drosophila* brain and to rule out contribution from potential sensory defects that would lead to inability to sense olfactory mating cues.

### The role of AMPK in regulation of sleep

Sleep is a highly conserved animal behavior, and humans spend roughly one-third of their life sleeping [63, 64]. Sleep and wakefulness are under the control of circadian and homeostatic processes. The circadian clock determines the timing and rhythmic nature of sleep onset, whereas homeostatic mechanisms are involved in sensing sleep drive and provide increasing sleep pressure as a function of time spent awake. Neurons have high demands for ATP, the major form of cellular energy, and low capacity to store nutrients [65], which has led to the hypothesis that sleep is required to replenish neuronal energy that is depleted during wakefulness [66]. This theory suggests that energy levels are reflected in glycogen and adenosine changes accumulated during metabolically demanding wakefulness and that these molecules play key roles in homeostatic sleep regulation [67]. Consistent with this notion, glycogen levels are indeed affected by the rest and wake cycle and drop after short periods of rest deprivation in *Drosophila* [68]. Adenosine is a breakdown product reflecting the depletion of ATP, the primary energy currency used by brain cells [69]. Consistent with the idea that sleep is necessary to reestablish energy stores, ATP has been shown in rats to increase during the initial hours of sleep when neuronal activity is low [31]. If the homeostat senses sleep drive by measuring energy levels, the molecular mechanism of the homeostat must involve an energy sensor that is activated by low cellular energy levels and initiates processes that restore energy levels to relieve sleep pressure after a nap. As the major cellular energy sensor activated by low energy levels, AMPK promotes processes that replenish energy levels [70]. Furthermore, the AMPKβ subunit contains a glycogen-binding domain that likely enables the AMPK complex to sense energy levels in the form of both ATP and glycogen in the nervous system [71]. If the AMPK complex is involved in homeostatic sleep regulation, then rebound sleep following sleep deprivation would not occur without this complex. Our results indicate that neuronal loss of AMPK affects sleep regulation and leads to loss of rebound sleep following deprivation, providing evidence that AMPK is indeed involved in homeostatic sleep regulation. Furthermore, our data indicates that AMPK is required for regulation of sleep in the central complex, the main structure of the brain has been associated with homeostatic sleep in *Drosophila* [37–40]. It will be of great interest to determine whether AMPK is part of the mechanism of sleep homeostasis and its role in psychiatric disorders characterized by sleep disturbances.

## Materials and methods

### Drosophila strains and maintenance

*Drosophila* larvae and adults of mixed sexes were raised on standard cornmeal medium (Nutri-Fly “Bloomington” formulation) under a 12:12 light/dark cycle at 25 °C with 60% humidity, unless otherwise stated. Fly lines *elav-GAL4; UAS-Dicer-2 (Dcr-2*) (#25750), *ppk-GAL4* (#32078), *UAS-mCD8::GFP* (#5137), *UAS-AMPKα-K56R* (*UAS-AMPKα-DN,* #50760), *ChAT-GAL4* (#6793), *189Y-GAL4* (#30817), *R24C10-GAL4* (#49075) [41, 42, 72], and *Canton S* were obtained from Bloomington *Drosophila* Stock Center (BDSC; Bloomington, IL). *UAS-alc-RNAi(KK*) (#109325) and the *w^1118^* (#60000) genetic background line were procured from Vienna *Drosophila* Resource Center (VDRC; Vienna, Austria). *UAS-alc-RNAi(8057-R2*) was obtained from the NIG-Fly stock center (Mishima, Shizuoka, Japan). The CRISPR/Cas9 system was used to create *alc::Venus* at the endogenous locus: sequence encoding a C-terminal Venus (yellow GFP derivative) tag was inserted upstream of the *alc* stop codon.

### Lifespan measurements

For starvation-survival experiments, thirty 3-5-day-old adult males were placed in vials containing 2% agar in water. Survival was assessed every 2-4 hours during the main course of the experiment for 9-10 replicate vials for each cross until all animals were dead. For each vial, the median survival was calculated in MatLab (The MathWorks, Inc., Natick, MA) as the time-point when the cumulative survivor function using the Kaplan-Meier method fell below 50%. For lifespan experiments, thirty male flies were collected upon eclosion into vials containing normal diet. Ten replicates for each cross were used, and the animals were transferred to fresh vials every 2 to 3 days. During each transfer, the numbers of dead animals left behind and carried over were recorded. Escaped flies or accidental deaths during transfer were recorded as censored. Longevity was monitored for 89 days. Survival data was analyzed for each vial in MatLab using the Kaplan-Meier nonparametric method accounting for censored data. As some control vials did not reach 50% mortality, to quantify survival, the Weibull distribution function was fit to the data, right-censoring the animals still alive at the end of the experiment. This analysis extends exponential distributions of failure (death) probability to allow for the increasing hazard rates associated with aging systems [73]. The scale parameter was determined for each vial and used for comparisons.

### Visualization of class-IV dendritic-arborization (da) neurons

Feeding and wandering third-instar larvae were selected and anesthetized by exposure to chloroform for 1 minute in a sealed container. Larvae were mounted in 90% glycerol, and GFP fluorescence in live animals was imaged. One individual GFP-labelled neuron, located in segment A2, was imaged per larva. Larvae were imaged at 20X using a Zeiss 780 LSM confocal microscope. Z-stacks were processed using FIJI (NIH) software [74], and analysis was performed using the TREES toolbox in MatLab [75].

### Courtship-conditioning assay

A courtship-conditioning assay was used to assay learning in adult flies[29]. Male flies were collected upon eclosion and housed individually until the start of the experiment. Virgin females were collected upon eclosion and housed in groups of 30. Mated females were generated by housing with males for 24 hours prior to experimental start, after which the males were removed. Male flies were split into two groups, naïve flies and flies to be trained. For training, individual male flies were incubated with mated females for 1 hour prior to testing in custom 2-cm-diameter mating chambers with a food source. For imaging courtship behavior, custom chambers were laser cut from clear acrylic. Each chamber set consisted of an array of 20, 1.5-cm individual chambers allowing for testing of all conditions in parallel (S3 Fig). Channels were cut to allow for a thin separator to be inserted through all of the chambers to keep the loaded males and female separate until the start of recording. Single naïve or trained males were then transferred into a testing arena together with individual virgin females (loaded through a separate loading hole and kept separate from males). Transfer was done using gentle aspiration to avoid disturbing the animals. Once all animals were loaded, the chamber was placed on a custom-built image-acquisition setup consisting of a 20-cm diffuse infrared LED backlight (Falcon illumination *FLFL-Si200-IR24*) and a Basler acA2000-50gmNIR GigE camera fitted with an IR filter (S3 Fig). Once the dividers separating males and females was removed, video was recorded for 10 minutes using LabView (National Instruments, Inc., Austin, TX). All experiments were done in a climate-controlled room at 25 °C and 70% humidity. Courtship behaviours were manually scored, with the scorer blinded to the genotype. The experiment was repeated 11 times, with three trials (individual fly pairs) per experiment. Courtship indices (CIs) were calculated as the percentage of time that a male fly spends courting during a 10-minute period. The learning index (LI) was calculated as the relative reduction of CI in trained male flies compared to naïve flies.

### Sleep assays and analysis

Locomotion over a 24-hour period was measured using the *Drosophila* Activity Monitor (DAM) system (TriKinetics, Inc., Waltham, MA). Adult males were collected in groups of 30 upon eclosion and housed under standard conditions until experimental start. Four-to-seven-day-old males were used for experiments, housed in 65-mm glass tubes with a plug of 5% sucrose and 2% agar medium at one end. Experiments were performed under a 12-hour light/12-hour dark cycle, and activity measurements were binned in one-minute periods. Episodes of sleep were defined as at least 5 minutes of uninterrupted quiescence. Animals with less than 10 minutes of activity during either the light or dark phase were flagged as dead. All analyses of sleep- and motion-bout dynamics were done in MatLab. For sleep-deprivation experiments, flies were mounted in DAM monitors and were mechanically stimulated using a vortexer mounting plate (TriKinetics) for 2 seconds every minute, over a 6-hour period prior to lights-on. Recovery sleep from flies with >60% loss of sleep during the deprivation period was analyzed and compared to baseline conditions 24 hours prior to the commencement of sleep deprivation. Recovery sleep was defined to occur in the first three hours following the end of sleep deprivation.

### Western Blotting

Fly heads were homogenized in Laemmli sample buffer (#1610737, Bio-Rad, Hercules, CA). For separation of polypeptides, samples were electrophoresed through precast polyacrylamide gels (Bio-Rad, #4561094) for 1 hour in a Tris/glycine/SDS buffer. Separated proteins were transferred to a nitrocellulose membrane and were then blocked with Odyssey blocking buffer (LI-COR Lincoln, NE, #927-40000) for 1 hour prior to incubation overnight with rabbit anti-pAMPKα (1:1000, Cell Signaling technology, Danvers, MA, #2535) and mouse anti-α-Tubulin (Sigma #T9026, diluted 1:5000) antibody at 4 °C. After 3 x 15-minute washes in PBS at RT, samples were incubated with IRDye 680RD and 800CW secondary antibodies diluted 1:10,000 (LI-COR) for 30 minutes at RT. Western blots were imaged using an Odyssey Fc imaging system (LI-COR) and ImageJ was used to quantify probe signal intensity.

### Immunostaining

Adult brains were dissected in cold Schneider’s medium and fixed in fresh 4% formaldehyde in PBS followed by four washes in PBST (PBS containing 3% Triton X-100). Brains were then blocked in PBST containing 3% normal goat serum (Sigma-Aldrich #G9023) for one hour at room temperature and incubated overnight at 4 °C with mouse anti-GFP (clone JL-8, #632381, Clontech) diluted 1:200 in PBST containing 3% normal goat serum. Tissues were then washed four times in PBST, incubated at room temperature with Alexa Fluor 488-labelled goat anti-mouse (Thermo Fisher Scientific, #A11001) and mounted in in ProLong Gold Antifade (Thermo Fisher). Imaging was performed using a Zeiss LSM 700 confocal microscope and Zen software, and image processing was done using the FIJI software package.

### Quantitative PCR andRNA sequencing

For quantitative real-time PCR (qPCR), total RNA was prepared from 10 adult male heads using the RNeasy Mini Kit (Qiagen #74106) with DNase treatment (Qiagen #79254). cDNA was synthesized using the iScript Reverse Transcription Supermix for RT-PCR (Bio-Rad #1708840), and qPCR was performed using the QuantiTect SYBR Green PCR Kit (Fisher Scientific #204145) on an Mx3005P qPCR system (Agilent Technologies). Expression was normalized to *RpL32.* Primers: *alc*; GGGCGACCATCAGTACAAGT and GCGTTCTCCACGCTTTTC; *RpL32,* AGTATCTGATGCCCAACATCG and CAATCTCCTTGCGCTTCTTG. For RNA-sequencing transcriptomics (RNA-seq), ten adult fly heads from 4-7-day-old males were harvested for each condition, and RNA libraries were prepared for Illumina HiSeq sequencing of paired-end 100-bp reads. Triplicates were sequenced for each genotype to determine differentially expressed genes.

### Mechanical stress test

The mechanical stress test was conducted on 3-day-old male flies as described [36]. For each genotype, three vials containing 10 flies each were evaluated, with genotypes masked during test and evaluation. Each vial of flies was subjected to 5 seconds of mechanical stimulation (vortexing), followed by 55 seconds of recovery. The number of upright flies was counted after this time and is represented as a percentage of total number of flies.

## Acknowledgements

We thank Annika Groenborg-Forsingdal for assistance with analysis of RNA-seq samples and Michael J. Texada for reading and editing the manuscript. This work was supported by the Danish Council for Independent Research, Medical Sciences grant 4183-00312 to K.F.R.

## Supporting information

**S1 Fig.**
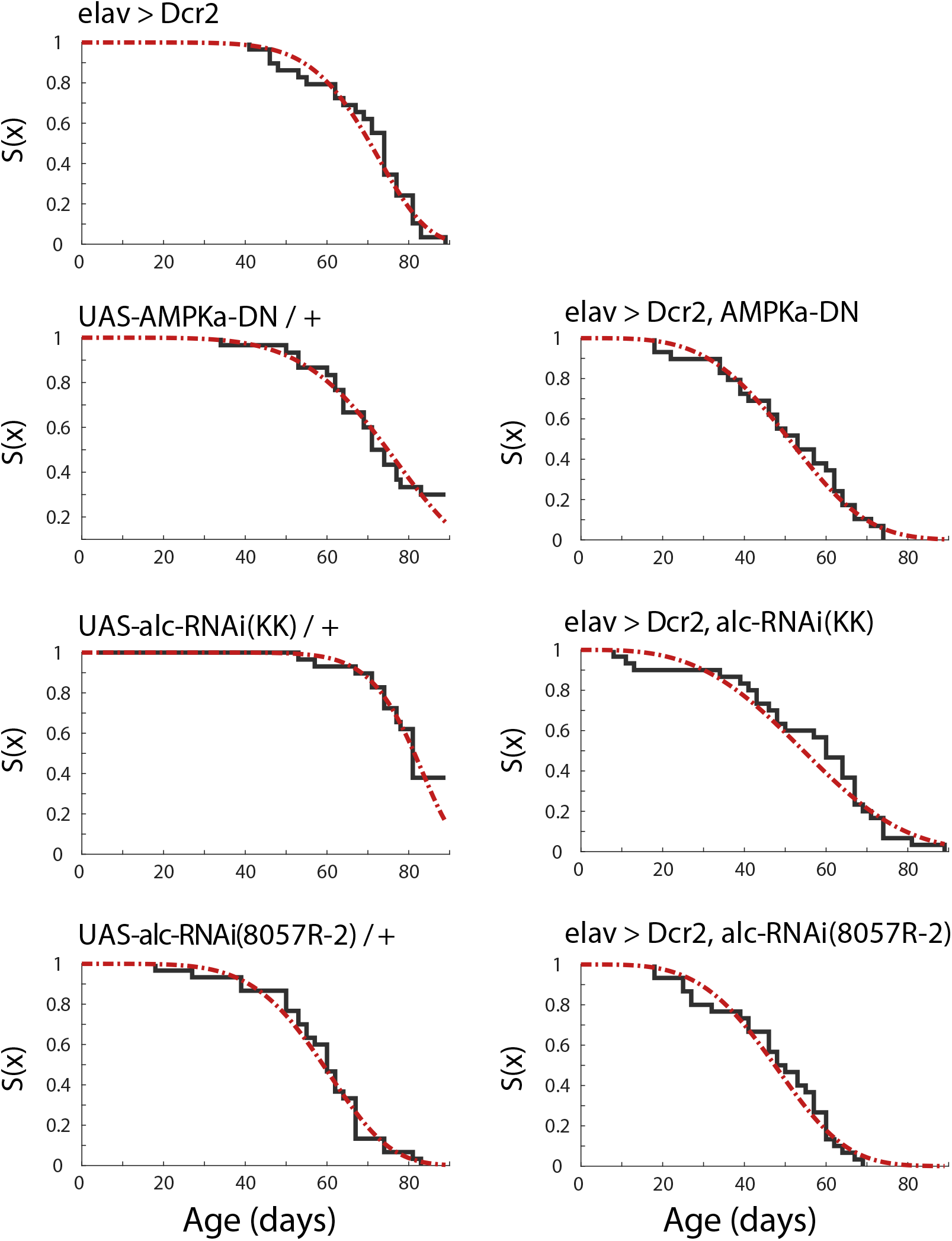
Example fits of survival data to a Weibull distribution. Example for all genotypes tested, showing fits of survival data, from individual vials of 30 animals, to a Weibull distribution.

**S2 Fig.**
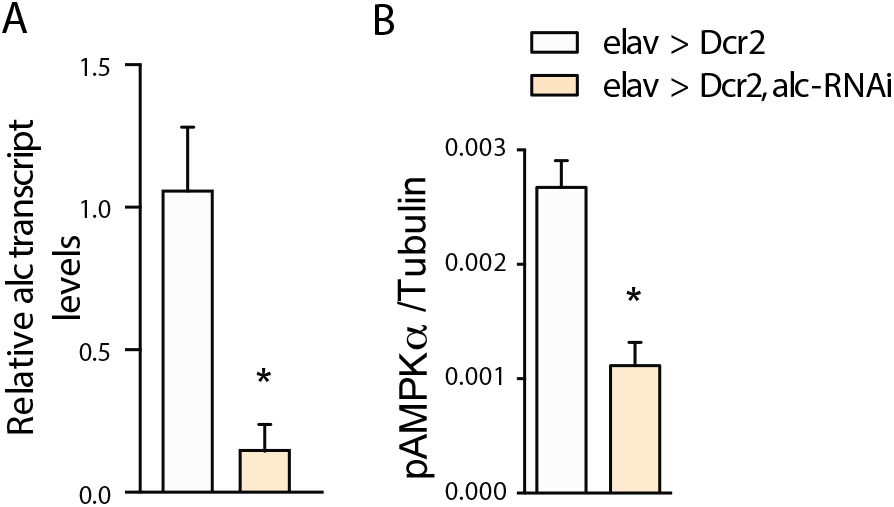
Pan-neuronal knockdown of *alc* reduces *alc* mRNA and levels of activated pAMPKα in adult heads. (A) Expression levels of *alc* mRNA in adult heads. Data shown is relative to driver control, set to 1.0. (B) Levels of pAMPKα are significantly reduced (−60% from controls) in adult heads when *alc* is knocked down pan-neuronally; *a* tubulin is used as a ratiometric loading control. Data from 4 experimental repeats using 8 adult heads per sample. Error bars indicate SEM. Mann Whitney test was used to determine statistical significance: *p<0.05, versus the control.

**S3 Fig.**
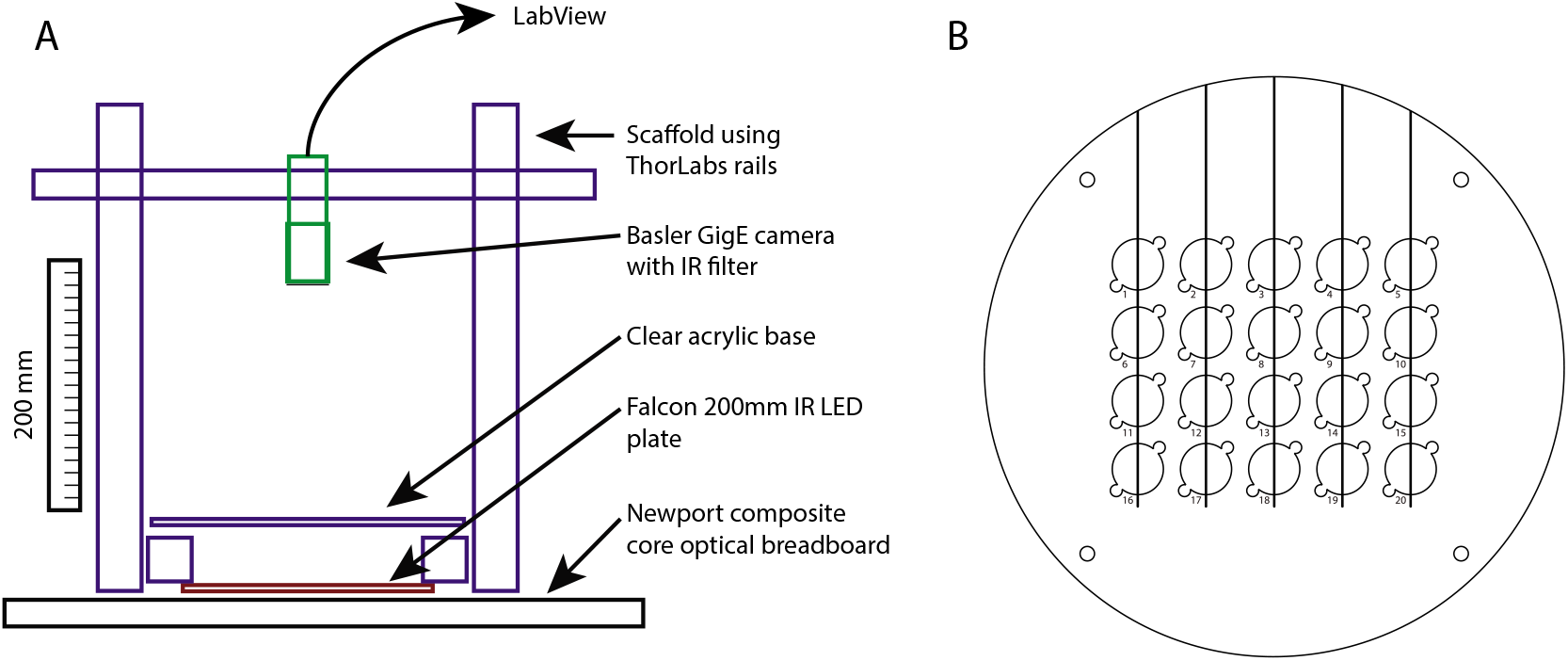
Schematic of courtship conditioning test setup. (A) Schematic diagram and components of the setup built to test courtship conditioning. (B) Schematic diagram of individual chambers laser-cut from acrylic that were used to test courtship conditioning.

**S4 Fig.**
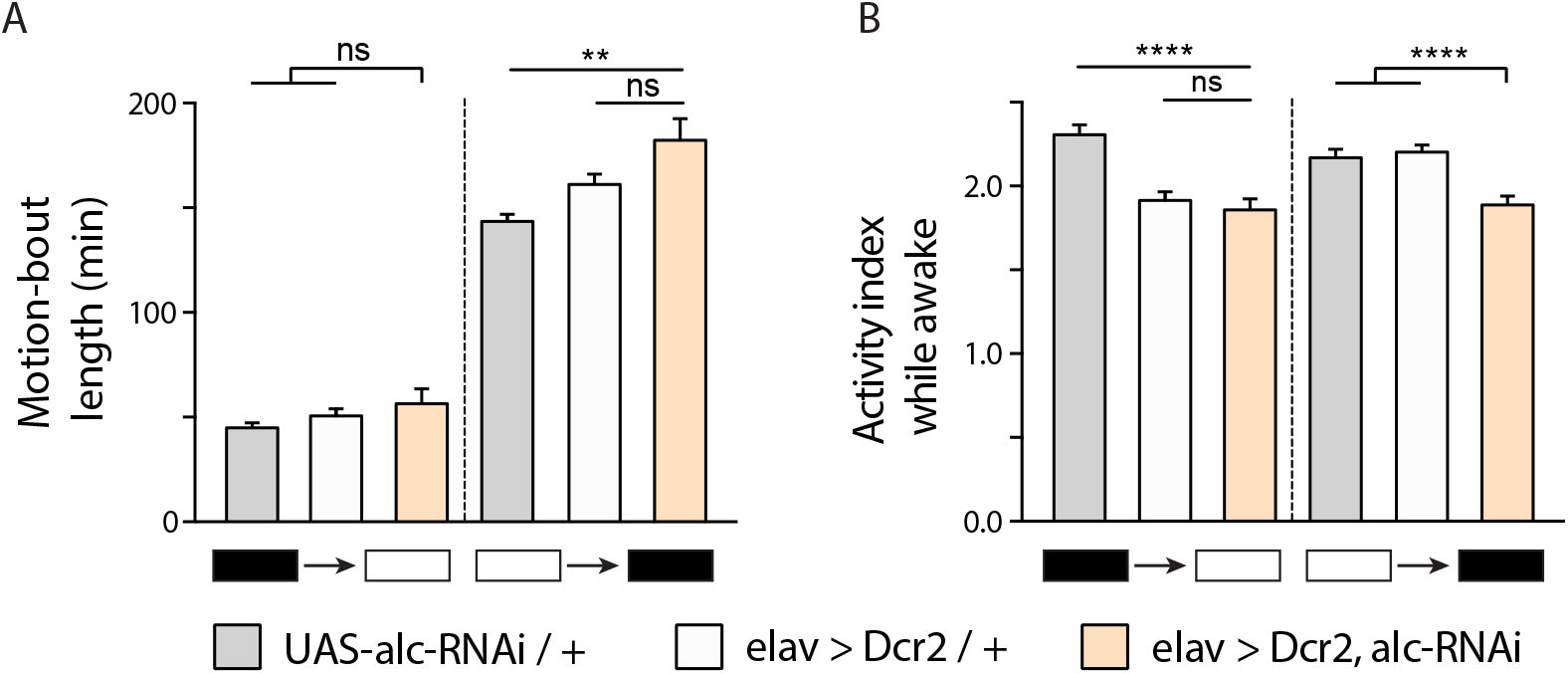
Knockdown of *alc/AMPKβ* in the nervous system does not increase motion-bout duration or levels of activity during light-dark transition periods. (A) Quantification of the average motion-bout length during dark-light and light-dark transitions (isolated as the motion bout present at t=0(24) and t=12 hours respectively) showed no significant differences in activity between *elav>Dcr-2, alc-RNAi* and control genotypes. (B) Quantification of activity (mean number of beam crossings per minute) during dark-light and light-dark transition periods showed a decrease in activity for *elav>Dcr-2, alc-RNAi* animals compared to control genotypes. Error bars indicate SEM. Kruskal-Wallis test with Dunn’s post-hoc testing was used to determine statistical significance: **p<0.01, ****p<0.0001, versus the control.

**S5 Fig.**
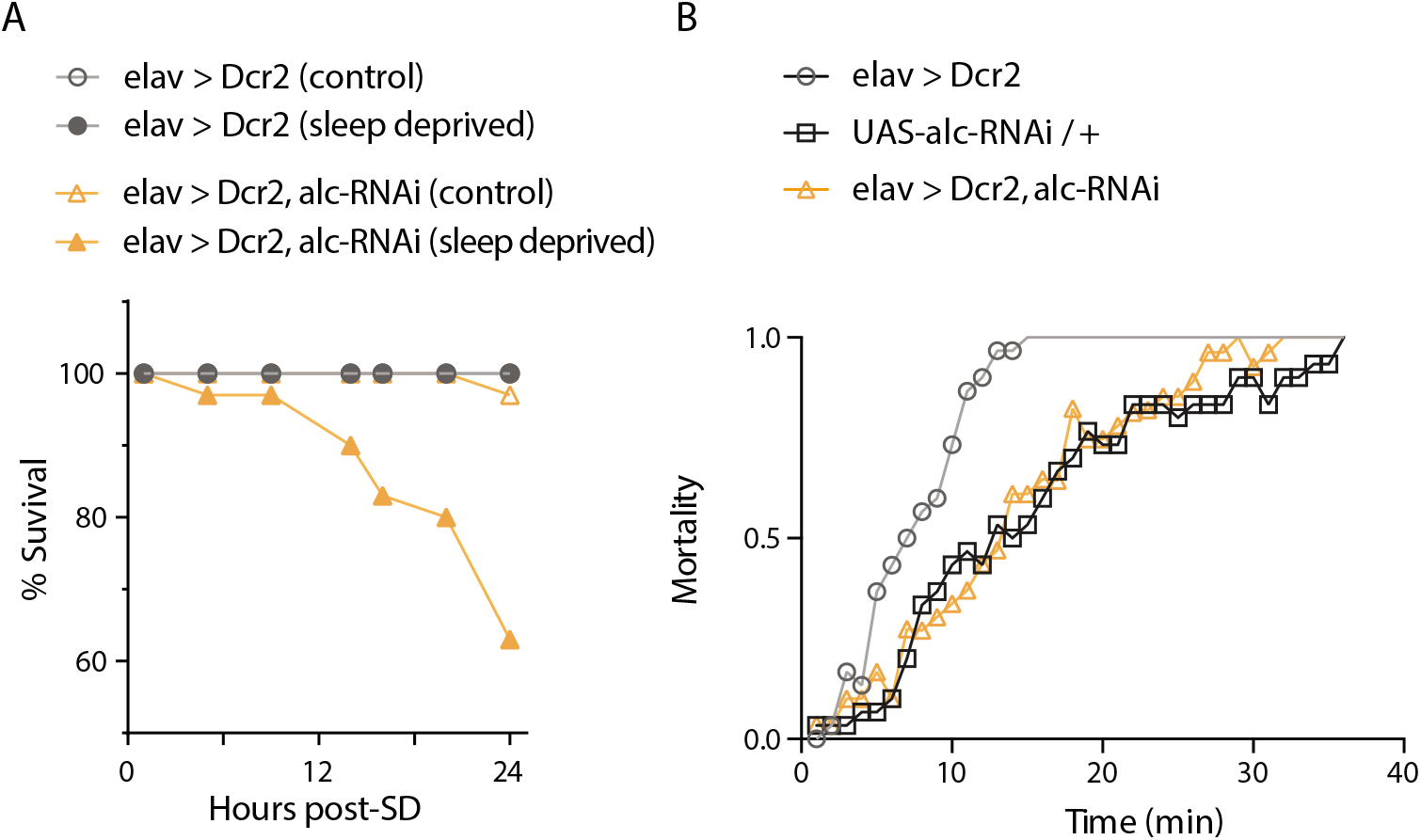
Knockdown of *alc/AMPKβ* in the nervous system increases mortality following sleep deprivation, but not mechanical stress. (A) Graph showing percentage survival and increased mortality of sleep deprived *elav>Dcr2, alc-RNAi* (n=32) animals, compared to a sleep-deprived (SD) control genotype (*elav>Dcr2/+,* n= 32), and non-sleep deprived controls (*elav>Dcr2/+, elav>Dcr2, alc-RNAi,* n=32). Survival was monitored over 24 hours post sleep deprivation, and the number of dead animals was counted. (B) Graph showing mean percentage survival following a mechanical-stress assay (n=3 vials of 10 animals for each genotype). Knockdown of *alc* in the nervous system does not increase susceptibility to mortality following mechanical stress.

**S6 Fig.**
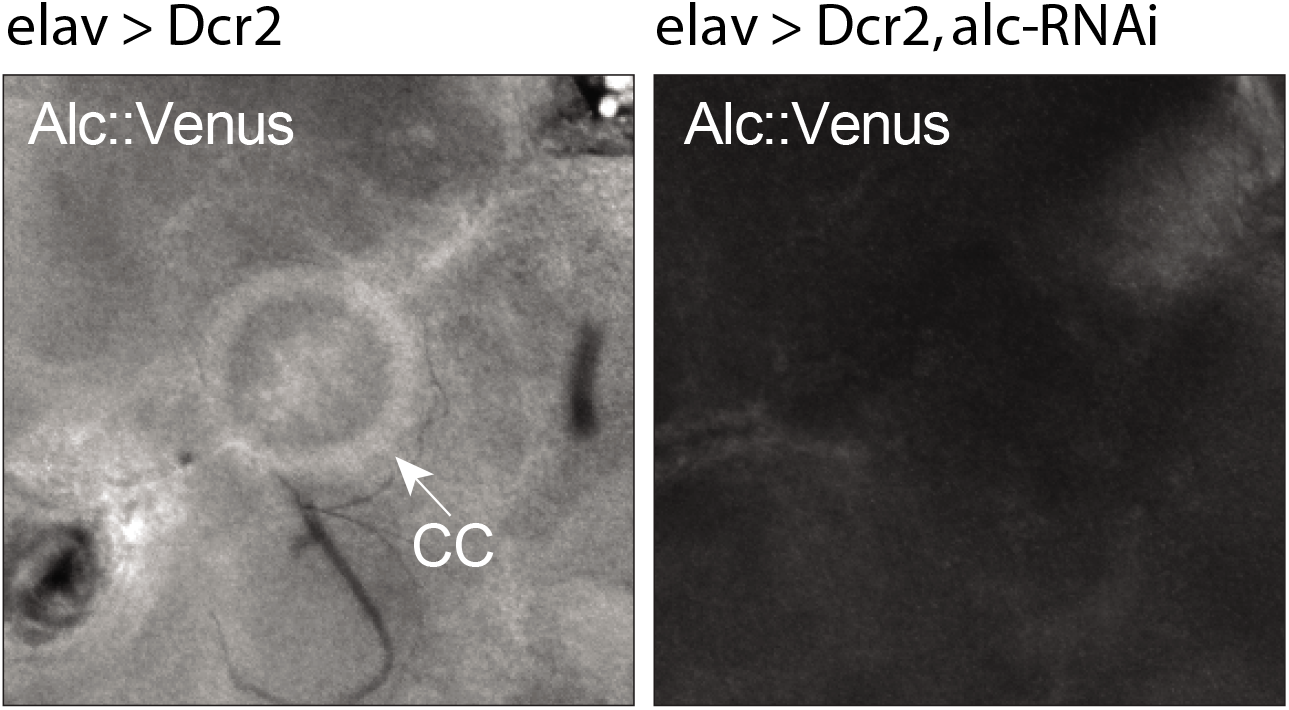
*alc/AMPKβ* is highly expressed in the central complex. Representative confocal image of adult brain showing concentration of Alc in the central-complex neuropil. Alc was imaged by using anti-GFP antibody to detect the expression of an endogenous Alc::Venus CRISPR knock-in. As expected, knockdown of *alc* in the nervous system greatly reduced detectible Alc::Venus in the brain.

**S1 Table.** RNA-seq data for differentially expressed genes in adult male fly heads between control and animals with knockdown of *alc* in the nervous system. Control *elav>Dcr2/+* animals were compared to *elav>Dcr2, alc-RNAi(KK*) animals with neuronal knockdown of *alc.*

